# *MLL4/KMT2D* mutations increase immune activity and predict therapy efficacy in colorectal cancer

**DOI:** 10.64898/2026.07.31.742076

**Authors:** Timothy En Haw Chan, H.T. Marc Timmers

## Abstract

**Background:** The *KMT2D* histone 3-lysine 4 methyltransferase (also known as MLL4) is a critical chromatin regulator, which is frequently inactivated by gene mutations in various types of cancer including colorectal cancer (CRC). Several lines of evidence suggest a strong association between chromatin regulation, tumor immunity and drug sensitivity. To explore the role of *KMT2D* in tumor immunity, we analyzed genomic and clinical datasets from The Cancer Genome Atlas and Memorial Sloan Kettering Cancer Center for various cancer entities.

**Result:** The results showed that *KMT2D*-mutated CRCs displayed a significantly higher expression of immune checkpoint regulators including *PD-L1*, *CTLA4* and *CD8* when compared to *KMT2D* wild-type CRCs. Mutations in *KMT2D* correlate with elevated T-effector and interferon-γ gene signatures indicating infiltration with active immune cells. By performing immune cell deconvolution from transcriptomic data, CRCs harboring *KMT2D* mutations are associated with increased infiltration of CD8+ T cells, NK cells and macrophages, but they display low amounts of Treg cells suggesting that KMT2D loss-of-function mutations correlate with an immunologically “hot tumor” phenotype. Furthermore, patients with *KMT2D* mutant CRC displayed better clinical responses to immune checkpoint inhibitor (ICI) therapy with an improved overall patient survival compared to patients with *KMT2D* wild-type CRC. Importantly, this effect did not exist in cohorts of CRC patients, which have not been treated with immunotherapies. To further understand the differential drug response effect related to *KMT2D*, we treated wild-type and *KMT2D* inactive epithelial cells with various cancer drugs. *KMT2D* mutant cells exhibited increased DNA damage and higher sensitivity to cisplatin in comparison to *KMT2D* wild-type cells.

**Conclusions:** *KMT2D* loss-of-function mutations are associated with a positive outcome of immunotherapy efficacy in CRC and responses to cisplatin treatments. The stratification of CRC patients by *KMT2D* gene status may enable personalized approaches by identifying CRC patient populations that benefit from ICI therapy and chemotherapy.

## Background

Colorectal cancer (CRC) represents a major global health burden as one of the most common malignancies in both incidence and mortality worldwide [1]. Although advances in cancer screening have improved disease outcomes, a substantial proportion of CRC patients still present with advanced-stage disease. Due to novel therapeutic approaches, the 5-year survival rate for metastatic CRC has increased to about 26% [2]. Among those novel therapeutic, immunotherapy is one of the most effective approaches in cancer treatment and the immune checkpoint inhibitors (ICIs) demonstrate significant efficacy across multiple cancer types [3,4]. While ICIs such as the PD-1 inhibitor pembrolizumab have shown promising results and gained FDA approval for advanced CRC with microsatellite instability or mismatch repair deficiency (MSI-H/dMMR), only a limited number of CRC patients harbor MSI-H/dMMR tumors and treatment resistance remains a clinical challenge [5–7]. There are four consensus molecular subtypes (CMSs) with distinguishing features identified in CRC including CMS1 which is characterized by hypermutations, MSI-H and strong immune activation predicting positive outcomes for immunotherapy [8,9]. However, alternative biomarkers such as tumor mutation burden (TMB) and PD-L1 expression remain inadequately characterized as predictors for immunotherapy outcomes in CRC [10]. Therefore, identification of novel biomarkers is essential to improve both patient stratification and therapeutic outcomes.

Emerging evidence underscores the critical importance of epigenetic mechanisms in mediating anti-tumor immunity [11]. Retrospective analyses in cancer show association of mutations in chromatin regulators with higher mutational burden and improved response to checkpoint immunotherapy including mutations in histone methyltransferase genes [12]. The KMT2 gene family constitutes a fundamental component of histone methyltransferases in human cells by regulating epigenetic pathways. *KMT2D* (also known as mixed-lineage leukemia protein 4 or *MLL4*) is frequently mutated in multiple cancer types including lung cancer and colorectal cancer [13,14]. *KMT2D* mutations occur mostly as frameshift-stop gain mutations and *KMT2D* mutant status has been correlated with diminished overall survival (OS) of non-small cell lung cancer patients [15]. Furthermore, pooled CRISPR mutagenic screening in a liver and gastric cancer mouse model identified *KMT2D* deficiency as a sensitizer to ICI [16,17]. Furthermore, the capacity of *KMT2D* to regulate integrin expression by T-cells [18], to activate dsRNA-interferon signaling [19] and to accelerate lymphomagenesis, which lead to increase anti-tumor immune responses [20,21]. Our own recent results in a human immortalized diploid epithelial cell line revealed that *KMT2D* is essential for the response to TGF-β, which is known to play an immuno-suppressive role in various cancers [22]. Taken together, these results suggest the potential of *KMT2D* to act as a biomarker predicting responses to ICIs.

In this study, we focused on the mutational characteristics of *KMT2D* in CRC to evaluate its alteration on immune activity and as a potential predictor for immunotherapy efficacy by assessing immune-related genes expression, immune infiltration cells, microsatellite-instability (MSI) status and Tumor Mutational Burden (TMB). Furthermore, we performed experimental work to study drug sensitivity of *KMT2D* inactivated normal diploid epithelial cells. Overall, this study indicates that *KMT2D* mutation status alone has a strong potential as a biomarker for immunotherapeutic efficacy and that *KMT2D* loss-of-function increases cisplatin sensitivity.

## Methods

### Data collection and differential expression genes analysis

Two public cohorts from the Cancer Genome Atlas (TCGA) and Memorial Sloan Kettering Cancer Center (MSKCC) IMPACT with tumor mutation status and genes expression information were used. To compare differential gene expression related to *KMT2D* mutation status, we used the Exploration function under Gene_Mutation module from TIMER2.0 [23] (https://compbio.top/timer2/) to check log2 fold changes of the differential expression of each gene for each cancer type. The Wilcoxon rank-sum test was used for comparison of the two groups.

### Immune cell infiltration analysis

To compare differential immune cell infiltration between tumors dependent on *KMT2D* mutation status, we used the Immune function under Mutation module from TIMER2.0 [23] (https://compbio.top/timer2/) to determine immune cell infiltration scores. TIMER and CIBERSOFT algorithms were used to determine the proportions of immune infiltrating cell types (IICs), based on deconvolution of immune gene expression signatures [24,25]. Default setting on TIMER2.0 database for TIMER and CIBERSOFT algorithms with the “Purity Adjustment” option and the Spearman’s correlation was used to perform the association analyses.

### MSI status, mutation count and tumor mutation burden

We downloaded MSI status, mutation count and tumor mutation burden data of TCGA from cBioPortal database (https://www.cbioportal.org). 406 samples from 401 patients in TCGA-COAD were grouped into two groups based on *KMT2D* mutation status to compare MSI status, mutation count and tumor mutation burden between the two groups. The Wilcoxon rank-sum test was used for comparison of the two groups.

### Overall Survival (OS) analysis

Two public cohorts from the TCGA-COAD (401 patients) and Memorial Sloan Kettering Cancer Center (MSKCC) IMPACT with ICI treatment (1662 patients) were used. Overall survival (OS) was analyzed by using the Kaplan-Meier plot option of cBioportal, in which two groups were separated by *KMT2D* mutation status of samples. The significance of OS was determined using the log-rank test.

### 3-(4,5-Dimethylthiazol-2-yl)-2,5-Diphenyltetrazolium Bromide (MTT) Cell Proliferation Assay

*KMT2D* wildtype and *KMT2D* CRISPR knockout hTERT RPE-1 cell lines were described in our previous study [22]. Cells were cultured in DMEM:F12 medium (Gibco) with 10% fetal bovine serum. 500 cells per well were seeded in 96-wells plates and incubated with different concentrations of cisplatin (0.4 – 50 μM), olaparin (0.4 – 50 μM) and doxorubicin (0.8 – 1000 μM) in triplicate. After 48 hours, the culture media was removed and cells were incubated with culture media including 2 mM 3-(4,5-dimethylthiazol-2-yl)-2,5-diphenyltetrazolium bromide (MTT) for 2 hours. After MTT incubation, 100 μl of DMSO were added after the MTT media were removed. The absorbence at 570 nm was determined on microplate reader (Tecan) after 15 min DMSO incubation. Overall cell viability results were calculated by GraphPad Prism 6.

## Results

### *KMT2D* genomic alterations lead to higher immune checkpoint genes expression in colorectal cancers

To investigate the involvement of *KMT2D* in CRC tumorigenesis, we performed expression analyses utilizing TCGA databases to evaluate *KMT2D* mRNA transcriptional activity of immune gene expression, *PD-L1(CD274), CTLA4, CD8A* and *CD8B* in tumor versus normal tissues. Among all cancer types covered by TCGA databases, colon adenocarcinomas (COAD), rectum adenocarcinomas (READ), skin cutaneous melanomas (SKCM) and uterine corpus endometrial carcinomas (UCEC) have significant overexpression of immune checkpoint genes compare with control tissues (Fig. 1A). *KMT2D* mutations were identified at allele frequencies exceeding 10% across distinct cancer types, including COAD (Fig. 1B). Due to the low case number in READ, we mostly focussed the analysis on COAD. This showed that *KMT2D* mutation in tumors correlates with significantly elevated *PD-L1* (*CD274)* (p=3.5×10^−8^), *CTLA4* (p=3.4×10^−7^), *CD8A* (p=3.5×10^−8^) and *CD8B* (p=0.004) mRNA expression in COAD (Fig. 1C). These association results suggest that *KMT2D* has a potential immune checkpoint regulator in colorectal cancers.

**Figure 1.**
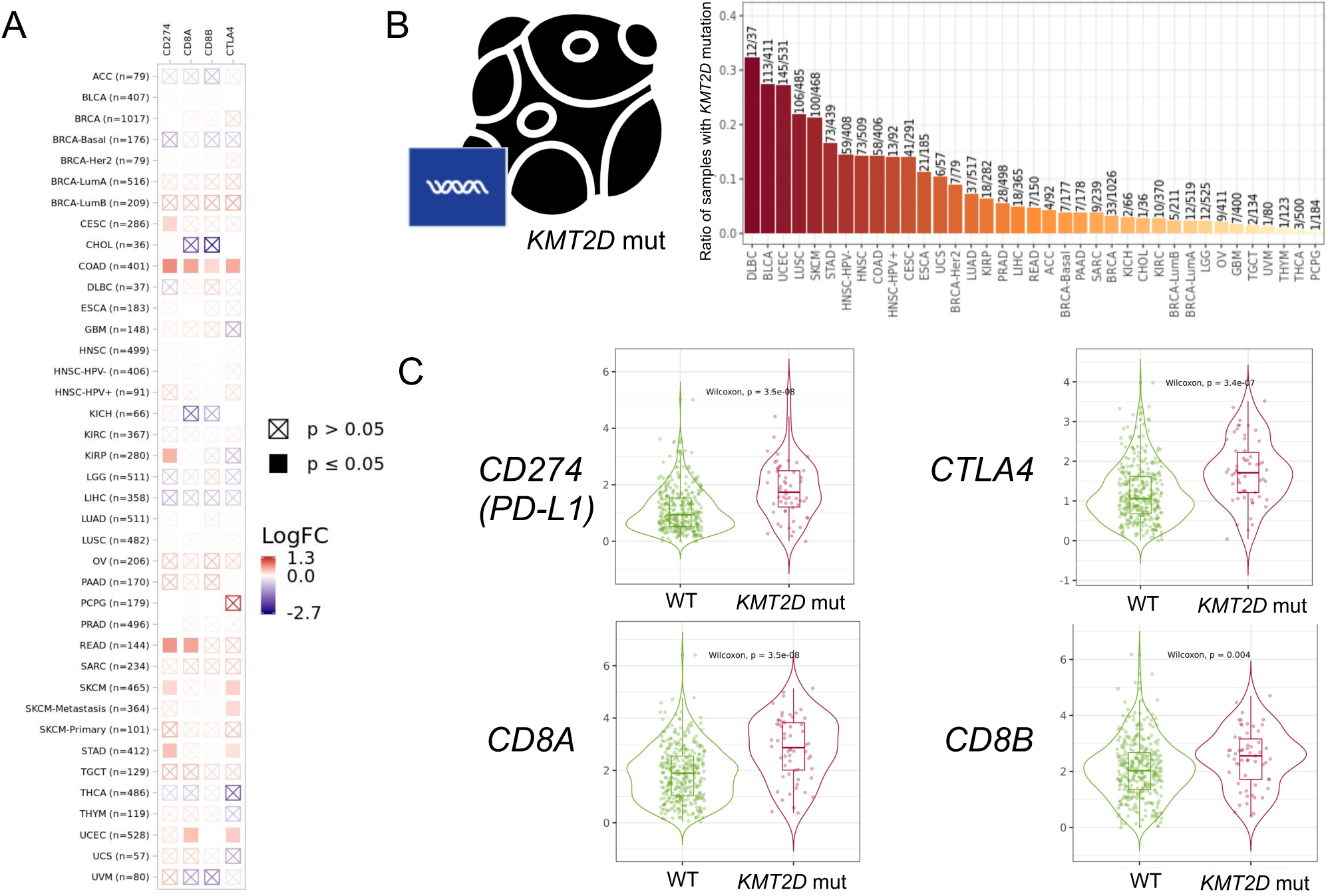
*KMT2D* mutation tumors have higher expression of *PD-L1, CTLA4* and *CD8* when compared to *KMT2D*-wild type CRC. (A) Relative expression level of *CD274, CTLA4, CD8A* and *CD8B* mRNAs in *KMT2D*-mutated tumors (n = 58) compared to wildtype *KMT2D-*tumors (n = 348) from TCGA database. (B) Proportion of tumors containing *KMT2D* mutation in different tumor types from TCGA. (C) Relative expression levels of *CD274, CTLA4, CD8A* and *CD8B* mRNAs in the TCGA-COAD dataset. Y-axis is relative mRNA expression level of tumor to normal in log2 fold changes.

### *KMT2D* mutation status reflects a higher immune activity tumor microenvironment in COAD

To elucidate the mechanistic basis linking *KMT2D* mutations to an increased immune cell activity, we assessed the impact of *KMT2D* status on T-effector and interferon gene signatures. The result showed a consistent positive correlation between *KMT2D* mutations and elevated T-effector and interferon genes, including in *CXCL10* (p=3.9×10^−8^), *CXCL9* (p=1.6×10^−10^), *GBP1* (p=2.1×10^−8^), *GZMB* (p=7.7×10^−4^), *IFI16* (p=2.4×10^−4^), *LAG3* (p=3.9×10^−12^), *OAS2* (p=2.4×10^−4^), *STAT1* (p=7.1×10^−6^) and *TAP1* (p=1.9×10^−4^) mRNAs level (Fig. 2). To understand differences in immune cell infiltration between *KMT2D*-WT *and KMT2D*-mut COAD tumors, we used CIBERSORT and TIMER 2.0 to perform a deconvolution analysis of the TCGA transcriptomic data [23–25]. This revealed that *KMT2D*-mut tumors are characterized by an enhanced infiltration of CD8+ T cells, NK T cell, macrophages and neutrophils, but by a decreased Treg cell infiltration (Fig. 3). Furthermore, in COAD *KMT2D* mutations correlated with MSI status (Chi-squared test, p=5.041×10^−5^), mutation count (Wilcoxon Test, p=2.878×10^−5^), and TMB (Wilcoxon Test, p=3.086×10^−5^) (Fig. 4). These observations collectively indicate that *KMT2D* mutation status associates with an immunologically active tumor microenvironment capable of enhanced ICI responsiveness.

**Figure 2.**
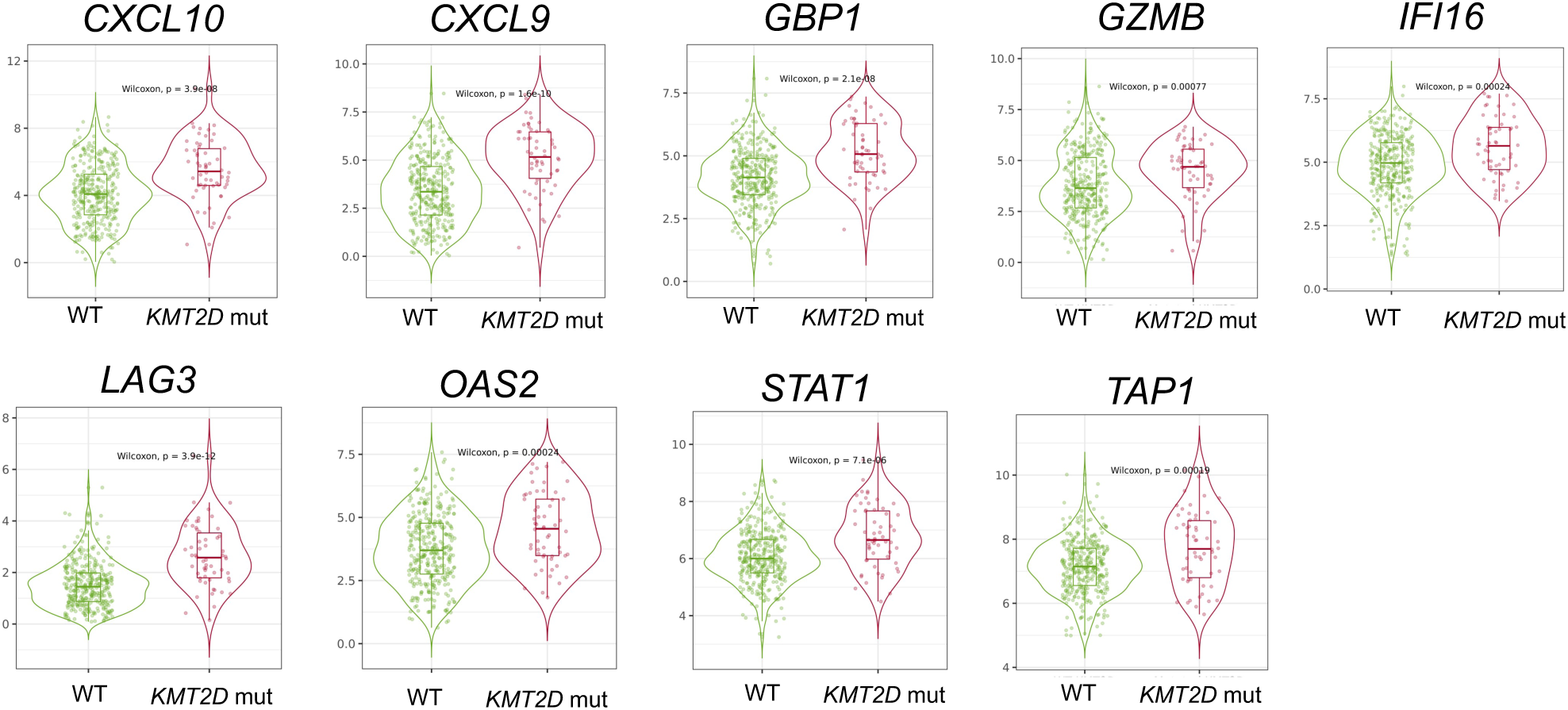
Association of *KMT2D* mutations with T-effector and interferon gene signatures in CRC. Relative expression level of the *CXCL10, CXCL9, GBP1, GZMB, IFI16, LAG3, OAS2, STAT1* and *TAP1* genes in *KMT2D* mutant and wildtype CRCs. Y-axis is relative mRNA expression level of tumor to normal in log2 fold changes.

**Figure 3.**
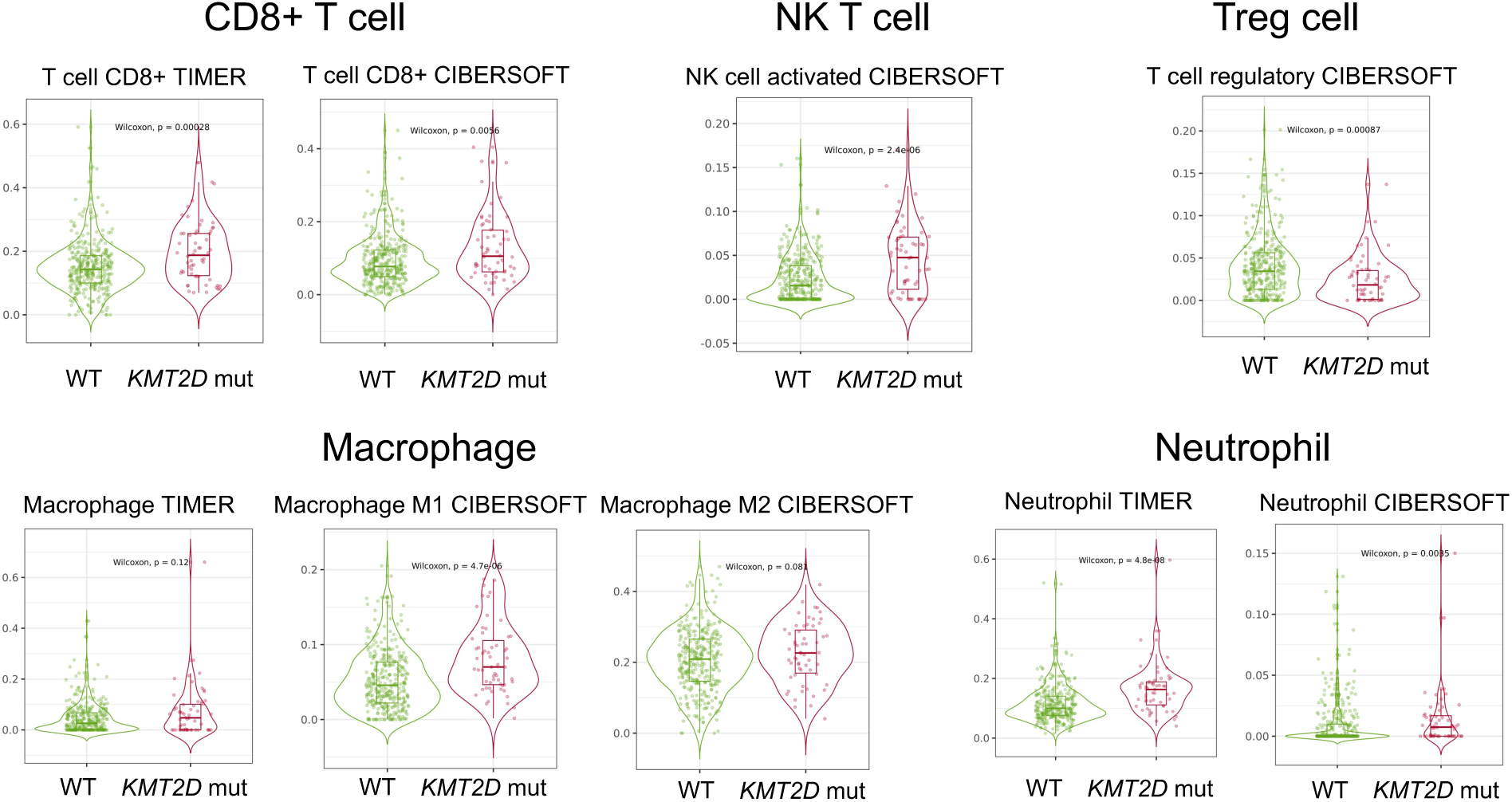
*KMT2D* mutations are associated with increase immune cell infiltration in CRC. Deconvolution of immune infiltrates of CD8+ T cells, NK T cells, Treg cells, macrophages and neutrophils from *KMT2D* mutant and *KMT2D* wildtype CRCs by TIMER and CIBERSOFT algorithms. Y-axis represents immune infiltration score and the significance of two group comparisons was done by the Wilcoxon Test and p-values are indicated.

**Figure 4.**
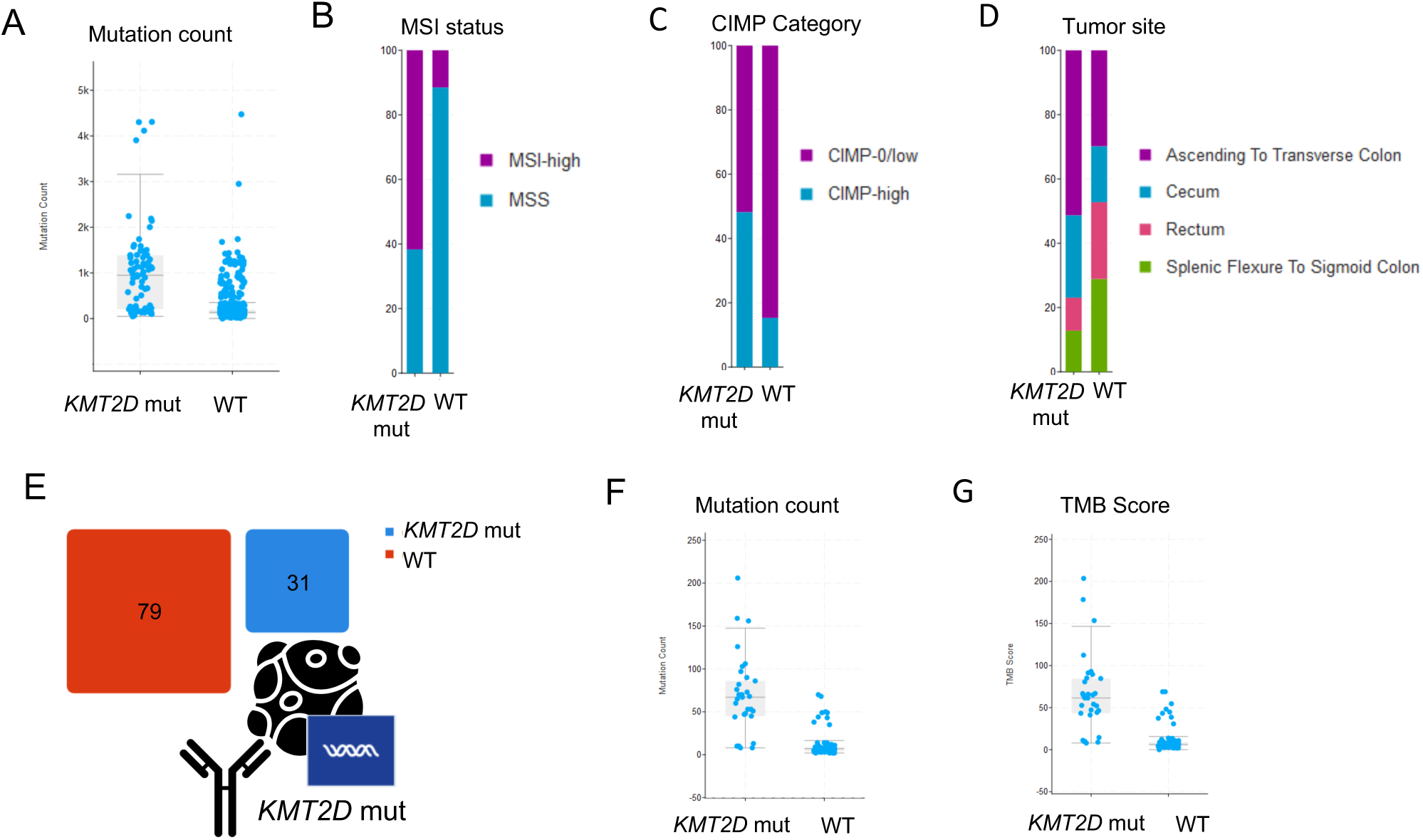
Associations of *KMT2D* mutations, MSI status, mutation count and tumor mutation burden in CRC. (A) Mutation count, (B) MSI status, (C) CIMP status and (D) tumor source from *KMT2D* mutant and *KMT2D* wildtype TCGA-COAD dataset. (E) Composition for MSK-IMPACT immune checkpoint inhibitor treatment colon cancer samples. (F) Mutation count and (G) TMB tumor mutation burden of MSK-IMPACT colon cancer samples dataset.

### *KMT2D* mutation correlate with ICI responses in CRC patients

To examine the correlation between *KMT2D* mutations and clinical outcome in CRC, we conducted an overall survival (OS) analysis within the TCGA-COAD cohort. The result show improved OS in patients with *KMT2D*-mut compared to *KMT2D*-WT, which did not reach significance (p = 0.535) (Fig. 5A). The TCGA-COAD cohort does not distinguish between ICI and non-ICI treated CRC patients. In contrast the MSKCC-IMPACT cohort, which includes only ICI treated patients, we found a significant improved OS in CRC patients with *KMT2D*-mut compared to *KMT2D*-WT (p = 0.01) (Fig. 5B). The findings suggest *KMT2D* status may constitute a valuable biomarker for CRC patients undergoing ICI therapy.

**Figure 5.**
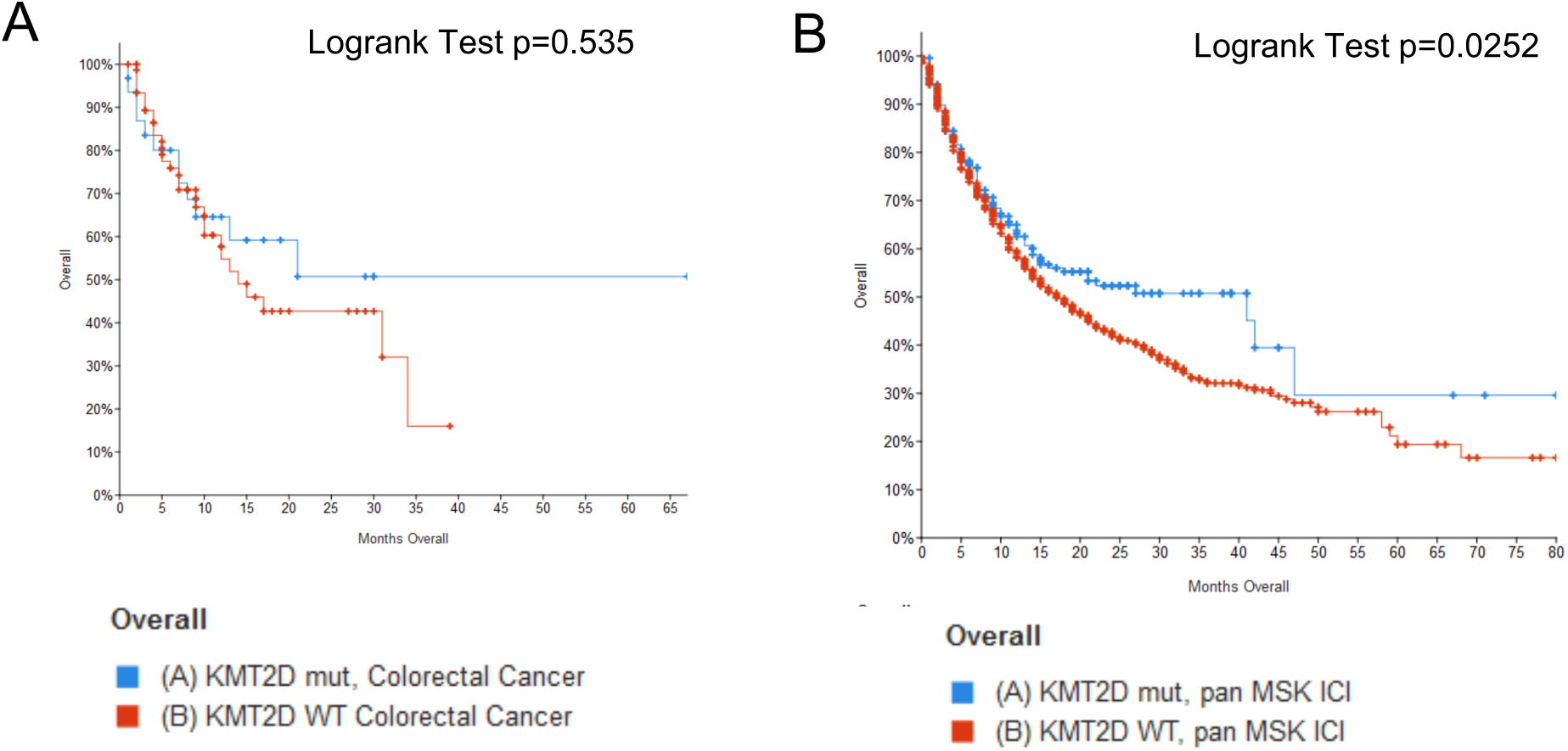
*KMT2D* mutations predict immune checkpoint therapy response. (A) Kaplan–Meier curves of overall survival grouping by *KMT2D* mutation status in the TCGA colon cancer cohort. (B) Kaplan–Meier curves of overall survival grouping by *KMT2D* mutation status in the MSKCC-IMPACT immune checkpoint inhibition cohort.

### *KMT2D* mutant cells display an increased cisplatin sensitivity

We recently discovered that *KMT2D* function is essential for an intact TGF-β response in a human immortalized epithelial cell line [22,26]. Besides its role as an immune cell suppressor [27,28], TGF-β is known to drive cisplatin resistance [29]. Therefore, we tested whether *KMT2D* inactivation sensitizes cells to cancer drugs inducing DNA damage. *KMT2D* wildtype and *KMT2D* knockout hTERT-immortalized diploid retinal pigment epithelial (RPE-1) cells were treated with different concentrations of the platinum-based anti-neoplastic agent cisplatin, the PARP inhibitor olaparin and the topoisomerase II inhibitor doxorubicin. Using the MTT assay for cell proliferation, we observed an increased sensitivity of cisplatin in *KMT2D* inactivated RPE-1 cells. The IC_50_ for cisplatin decreased from 21.5 µM for *KMT2D* wildtype to 8.1 µM for *KMT2D* mutant cells (Fig. 6A). No increased sensitivity was observed for olaparin and doxorubicin, which may indicate that the single-strand DNA breaks and double-strand DNA break repair system are normally functional in *KMT2D* inactivated cells (Fig. 6B,C). In contrast, efficient removal of cisplatin-induced DNA crosslinks seems to require wild type KMT2D.

**Figure 6.**
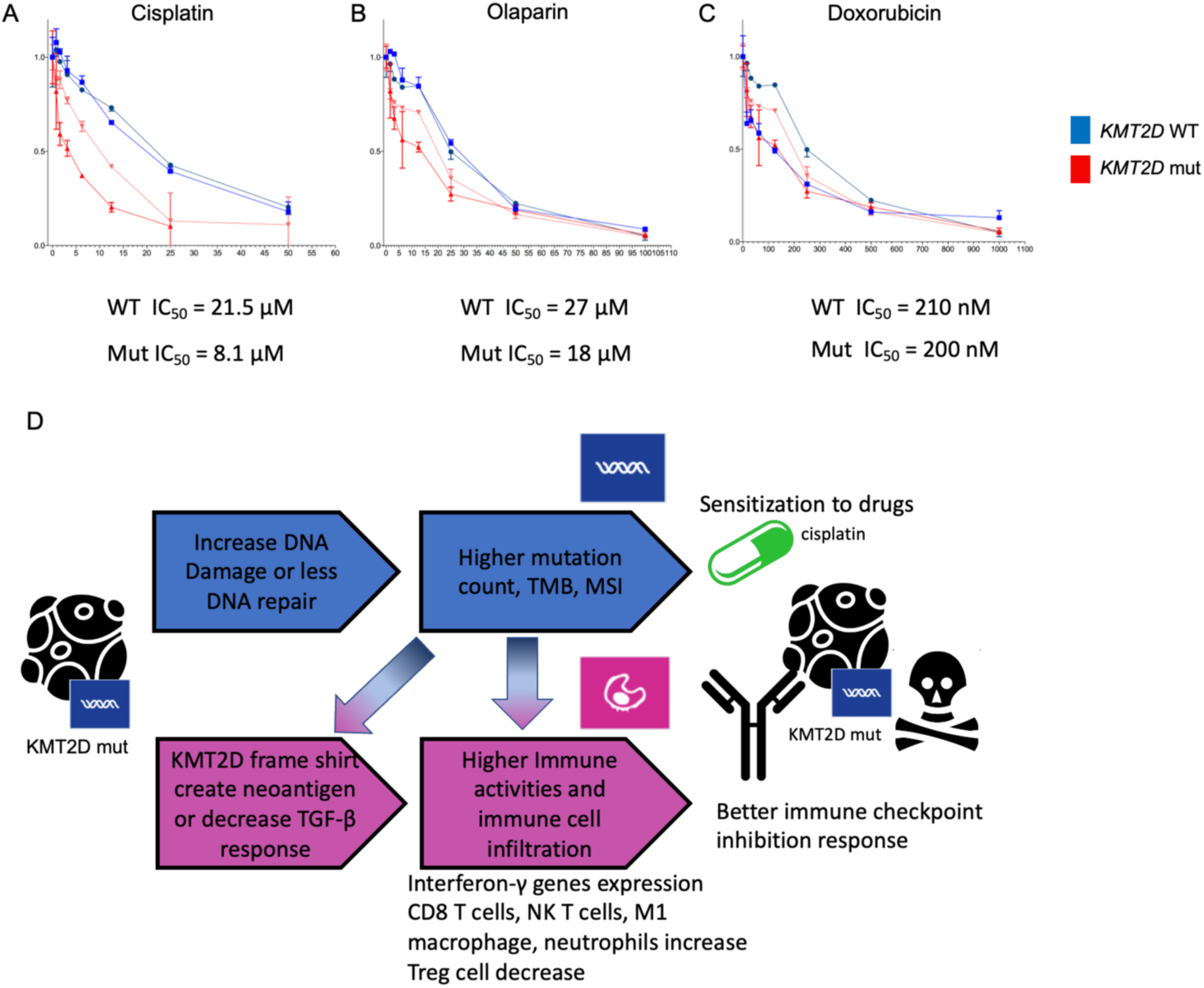
*KMT2D* mutant cells increase cisplatin sensitivity and possible mechanism of *KMT2D* mutant cells thought increase DNA damage response and treatment sensitivity. (A) IC_50_ of (A) cisplatin, (B) PARP inhibitor olaparin and (C) doxorubicin for wild type (WT) and *KMT2D* mutant (Mut) cells. (D) A hypothetical working model for the mechanism of *KMT2D* mutant tumor in enhance immune activity and different drug responses.

## Discussion

Recent years have witnessed improvements in immune checkpoint inhibitor (ICI) efficacies for diverse cancer types [30,3,4]. In this study, we found that *KMT2D* mutations may act as a favorable prognostic indicator for improved clinical outcomes in colorectal carcinoma (CRC) patients receiving ICI therapy. We employed bioinformatics approaches to explore potential mechanisms of *KMT2D*-mediated immunotherapy enhancement in CRC, which were previously investigated in mouse liver cancer and lymphoma models [16,19,21]. Our study is a retrospective analysis relying on TCGA and MSK-IMPACT public cohorts subjected to whole exome sequencing or targeted panel sequencing. In this we uncovered associations between *KMT2D* mutations and tumor-infiltrating immune activity. We proposed that *KMT2D* status may act as predictor of immunotherapy responses independent of other biomarkers such as microsatellite-instability–high (MSI-H), mismatch-repair–deficient (dMMR) or DNA polymerase epsilon (POLE) mutations [31,32]. Interestingly, CRC patients with *KMT2D* mutation tumors, who are receiving conventional therapies, did not demonstrate an equivalent clinical benefit in terms of OS. Exploratory analyses implicate elevated tumor burden (TMB) and microsatellite instability (MSI) scores as potential mechanistic mediators of *KMT2D*-driven therapeutic benefit in CRC. CIBERSORT and TIMER deconvolution approaches revealed substantially increased populations of CD8+ T cells, macrophages and neutrophils with decreased Treg cell abundances in *KMT2D*-mutant tumors suggesting that *KMT2D* gene status may represent a positive predictor of ICI efficacy in CRC patients.

KMT2D gained attention as a prominent epigenetic regulatory protein, which links the transcriptional activation of genes via histone H3-lysine 4 methylation and to the recruitment of other epigenetic regulators [33,34]. Comprehensive cancer genome studies have identified *KMT2D* as one of the most frequently altered genes across diverse cancers [35,36]. *KMT2D* mutations have been demonstrated to facilitate epithelial-to-mesenchymal plasticity [37,38]. Additionally, *KMT2D* participates in maintaining genomic integrity as *KMT2D* mutations result in a substantial elevation in (TMB) across multiple cancer types [39]. Recent studies show the alterations of *KMT2C*, which is paralogous to *KMT2D*, improved immunotherapy responses [40–43]. While a previous study reported *KMT2D* exon 39 mutations and combination of *KMT2C* and *KMT2D* mutation group correlate with a more abundant immune cell infiltration in CRC [44,45], our investigations in CRC further validate a consistent positive relationship between mutations across a whole *KMT2D* gene alone with an elevated TMB, which may translate to an increased neo-antigen expression. Elevated neo-antigen load could lead to recognition of tumor cells by cytotoxic T cells and macrophages facilitating anti-tumor immune responses, cisplatin responses and improved clinical outcomes. Recently, we discovered *KMT2D* is required for efficient TGF-β responses via a feed-forward loop with the JUNB transcription factor [22,26]. As TGF-β is a well-known effector of immune evasion [46,47] and drives cisplatin resistance [29], we speculate that loss of TGF-β responses contributes to immune activity and cisplatin sensitivity of *KMT2D* mutant tumors.

## Conclusion

Collectively, our association analyses and proposed molecular mechanism provide rationale for the connection of *KMT2D* mutations with improved ICI treatment outcomes. The stratification of CRC patients by *KMT2D* gene status as biomarker may enable personalized approaches by identifying CRC patient populations that benefit from ICI therapy. Our results are based on retrospective analyses of the TCGA dataset and they provide directions for future validations and investigations to elucidate the precise immunological mechanisms underlying *KMT2D*-driven immune modulation and drug sensitivity.

## List of abbreviations

COAD: colon adenocarcinoma
CRC: colorectal cancer
CMSs: consensus molecular subtypes
ICI: immune checkpoint inhibitor
IICs: immune infiltrating cell types
MSI: microsatellite-instability
MSI-H: microsatellite-instability–high
dMMR: mismatch-repair–deficient
MLL4: mixed-lineage leukemia protein 4
MSKCC: Memorial Sloan Kettering Cancer Center
MTT: 3-(4,5-dimethylthiazol-2-yl)-2,5-diphenyltetrazolium bromide
OS: overall survival ()
POLE: polymerase epsilon
READ: rectum adenocarcinoma
TCGA: the Cancer Genome Atlas
TMB: tumor mutation burden
UCEC: uterine corpus endometrial carcinoma
SKCM: skin cutaneous melanoma

## Declarations

### Ethics approval and consent to participate

Not applicable

### Consent for Publication

Not applicable

### Availability of Data and Materials

The datasets analyzed for this study can be found in the cBioPortal database (https://www.cbioportal.org) and TIMER 2.0 database (http://cistrome.org/TIMER, https://compbio.top/timer2/).

### Author Contributions

TEHC: Conceptualization. Data curation, Analysis, Investigation, Experiment, Writing – original draft. HTMT: Conceptualization, Funding acquisition, Supervision, Writing – review and editing.

## Acknowledgments

We thank all Timmers lab members for their input and constructive discussions and especially Sheikh Nizamuddin for bioinformatic assistance and critical reading of this manuscript. We also thank Hui-Ru Chen (Hospital for Special Surgery, NY) for critical reading of the manuscript.

## Reference

1. The Cancer Genome Atlas Network. Comprehensive molecular characterization of human colon and rectal cancer. Nature. 2012;487:330–7. 10.1038/nature11252

2. Zeineddine FA, Zeineddine MA, Yousef A, Gu Y, Chowdhury S, Dasari A, et al. Survival improvement for patients with metastatic colorectal cancer over twenty years. npj Precis Onc. 2023;7:16. 10.1038/s41698-023-00353-4

3. He X, Xu C. Immune checkpoint signaling and cancer immunotherapy. Cell Res. 2020;30:660–9. 10.1038/s41422-020-0343-4

4. Sharma P, Goswami S, Raychaudhuri D, Siddiqui BA, Singh P, Nagarajan A, et al. Immune checkpoint therapy—current perspectives and future directions. Cell. 2023;186:1652–69. 10.1016/j.cell.2023.03.006

5. Le DT, Uram JN, Wang H, Bartlett BR, Kemberling H, Eyring AD, et al. PD-1 Blockade in Tumors with Mismatch-Repair Deficiency. N Engl J Med. 2015;372:2509–20. 10.1056/NEJMoa1500596

6. Le DT, Durham JN, Smith KN, Wang H, Bartlett BR, Aulakh LK, et al. Mismatch repair deficiency predicts response of solid tumors to PD-1 blockade. Science. 2017;357:409–13. 10.1126/science.aan6733

7. Samstein RM, Lee C-H, Shoushtari AN, Hellmann MD, Shen R, Janjigian YY, et al. Tumor mutational load predicts survival after immunotherapy across multiple cancer types. Nat Genet. 2019;51:202–6. 10.1038/s41588-018-0312-8

8. Guinney J, Dienstmann R, Wang X, De Reyniès A, Schlicker A, Soneson C, et al. The consensus molecular subtypes of colorectal cancer. Nat Med. 2015;21:1350–6. 10.1038/nm.3967

9. Dienstmann R, Vermeulen L, Guinney J, Kopetz S, Tejpar S, Tabernero J. Consensus molecular subtypes and the evolution of precision medicine in colorectal cancer. Nat Rev Cancer. 2017;17:79–92. 10.1038/nrc.2016.126

10. Yarchoan M, Albacker LA, Hopkins AC, Montesion M, Murugesan K, Vithayathil TT, et al. PD-L1 expression and tumor mutational burden are independent biomarkers in most cancers. JCI Insight. 2019;4:e126908. 10.1172/jci.insight.126908

11. Cao J, Yan Q. Cancer Epigenetics, Tumor Immunity, and Immunotherapy. Trends in Cancer. 2020;6:580–92. 10.1016/j.trecan.2020.02.003

12. Gjorgjievska M, Bukovec D, Mehandziska S, Risteski M, Kungulovski I, Mitrev Z, et al. Tumors with mutations in chromatin regulators are associated with higher mutational burden and improved response to checkpoint immunotherapy. Clin Epigenet. 2025;18:19. 10.1186/s13148-025-02038-0

13. Guo C, Chen LH, Huang Y, Chang C-C, Wang P, Pirozzi CJ, et al. KMT2D maintains neoplastic cell proliferation and global histone H3 lysine 4 monomethylation. Oncotarget. 2013;4:2144–53. 10.18632/oncotarget.1555

14. Alam H, Tang M, Maitituoheti M, Dhar SS, Kumar M, Han CY, et al. KMT2D Deficiency Impairs Super-Enhancers to Confer a Glycolytic Vulnerability in Lung Cancer. Cancer Cell. 2020;37:599–617.e7. 10.1016/j.ccell.2020.03.005

15. Shi Y, Lei Y, Liu L, Zhang S, Wang W, Zhao J, et al. Integration of comprehensive genomic profiling, tumor mutational burden, and PD-L1 expression to identify novel biomarkers of immunotherapy in non-small cell lung cancer. Cancer Medicine. 2021;10:2216–31. 10.1002/cam4.3649

16. Wang G, Chow RD, Zhu L, Bai Z, Ye L, Zhang F, et al. CRISPR-GEMM Pooled Mutagenic Screening Identifies KMT2D as a Major Modulator of Immune Checkpoint Blockade. Cancer Discovery. 2020;10:1912–33. 10.1158/2159-8290.CD-19-1448

17. Wang N, Li D, Zhang T, Pachai MR, Schoeps DM, Bao Y, et al. Loss of Kmt2c/d promotes gastric cancer and confers vulnerability to mTORC1 and PD-1 inhibition. Journal of Clinical Investigation. 2026;136:e194462. 10.1172/JCI194462

18. Potter SJ, Zhang L, Kotliar M, Wu Y, Schafer C, Stefan K, et al. KMT2D regulates activation, localization, and integrin expression by T-cells. Frontiers in immunology. 2024;15:1341745. 10.3389/fimmu.2024.1341745

19. Ning H, Huang S, Lei Y, Zhi R, Yan H, Jin J, et al. Enhancer decommissioning by MLL4 ablation elicits dsRNA-interferon signaling and GSDMD-mediated pyroptosis to potentiate anti-tumor immunity. Nature Communications. 2022;13:6578. 10.1038/s41467-022-34253-1

20. Vlasevska S, Garcia-Ibanez L, Duval R, Holmes AB, Jahan R, Cai B, et al. KMT2D acetylation by CREBBP reveals a cooperative functional interaction at enhancers in normal and malignant germinal center B cells. Proc Natl Acad Sci USA. 2023;120:e2218330120. 10.1073/pnas.2218330120

21. Li J, Chin CR, Ying H-Y, Meydan C, Teater MR, Xia M, et al. Loss of CREBBP and KMT2D cooperate to accelerate lymphomagenesis and shape the lymphoma immune microenvironment. Nature Communications. 2024;15:2879. 10.1038/s41467-024-47012-1

22. Chan TEH, Islam MS, Fotouhi O, Bozkurt M, Nizamuddin S, Schüle KM, et al. MLL4/KMT2D histone methyltransferase and JUNB cooperate in a feed-forward loop to support AP-1-dependent TGF-β signaling. Genes Dev. 2026;40:1012–28. 10.1101/gad.353313.125

23. Li T, Fu J, Zeng Z, Cohen D, Li J, Chen Q, et al. TIMER2.0 for analysis of tumor-infiltrating immune cells. Nucleic Acids Research. 2020;48:W509–14. 10.1093/nar/gkaa407

24. Newman AM, Steen CB, Liu CL, Gentles AJ, Chaudhuri AA, Scherer F, et al. Determining cell type abundance and expression from bulk tissues with digital cytometry. Nat Biotechnol. 2019;37:773–82. 10.1038/s41587-019-0114-2

25. Newman AM, Liu CL, Green MR, Gentles AJ, Feng W, Xu Y, et al. Robust enumeration of cell subsets from tissue expression profiles. Nat Methods. 2015;12:453–7. 10.1038/nmeth.3337

26. Baas R, van Teeffelen H, Tjalsma SJD, Timmers HTM. The mixed lineage leukemia 4 (MLL4) methyltransferase complex is involved in transforming growth factor beta (TGF-beta)-activated gene transcription. Transcription. 2018;9:67–74. 10.1080/21541264.2017.1373890

27. Batlle E, Massagué J. Transforming Growth Factor-β Signaling in Immunity and Cancer. Immunity. 2019;50:924–40. 10.1016/j.immuni.2019.03.024

28. Liu M, Kuo F, Capistrano KJ, Kang D, Nixon BG, Shi W, et al. TGF-β suppresses type 2 immunity to cancer. Nature. 2020;587:115–20. 10.1038/s41586-020-2836-1

29. Oshimori N, Oristian D, Fuchs E. TGF-β Promotes Heterogeneity and Drug Resistance in Squamous Cell Carcinoma. Cell. 2015;160:963–76. 10.1016/j.cell.2015.01.043

30. Pardoll DM. The blockade of immune checkpoints in cancer immunotherapy. Nat Rev Cancer. 2012;12:252–64. 10.1038/nrc3239

31. André T, Shiu K-K, Kim TW, Jensen BV, Jensen LH, Punt C, et al. Pembrolizumab in Microsatellite-Instability–High Advanced Colorectal Cancer. N Engl J Med. 2020;383:2207–18. 10.1056/NEJMoa2017699

32. Maddalena G, Zeineddine FA, Chowdhury S, Zeineddine MA, Yousef AM, Bergamo F, et al. Prognosis and treatment response stratification according to loss of proofreading (LOP) *POLE* variants. J Immunother Cancer. 2025;13:e012190. 10.1136/jitc-2025-012190

33. Froimchuk E, Jang Y, Ge K. Histone H3 lysine 4 methyltransferase KMT2D. Gene. 2017;627:337–42. 10.1016/j.gene.2017.06.056

34. Wang L-H, Aberin MAE, Wu S, Wang S-P. The MLL3/4 H3K4 methyltransferase complex in establishing an active enhancer landscape. Biochemical Society Transactions. 2021;49:1041–54. 10.1042/BST20191164

35. Fagan RJ, Dingwall AK. COMPASS Ascending: Emerging clues regarding the roles of MLL3/KMT2C and MLL2/KMT2D proteins in cancer. Cancer Letters. 2019;458:56–65. 10.1016/j.canlet.2019.05.024

36. Zhang Y, Lv K, Ma X, Wang L, Xu Y. The MLL4: Roles in cell differentiation, adipogenesis and cancer. Biochemical Pharmacology. 2025;242:117371. 10.1016/j.bcp.2025.117371

37. Zhang Y, Donaher JL, Das S, Li X, Reinhardt F, Krall JA, et al. Genome-wide CRISPR screen identifies PRC2 and KMT2D-COMPASS as regulators of distinct EMT trajectories that contribute differentially to metastasis. Nature Cell Biology. Nature Publishing Group; 2022;24:554–64. 10.1038/s41556-022-00877-0

38. Seehawer M, Li Z, Nishida J, Foidart P, Reiter AH, Rojas-Jimenez E, et al. Loss of Kmt2c or Kmt2d drives brain metastasis via KDM6A-dependent upregulation of MMP3. Nature Cell Biology. 2024;26:1165–75. 10.1038/s41556-024-01446-3

39. Kantidakis T, Saponaro M, Mitter R, Horswell S, Kranz A, Boeing S, et al. Mutation of cancer driver *MLL2* results in transcription stress and genome instability. Genes Dev. 2016;30:408–20. 10.1101/gad.275453.115

40. Jiao Y, Lv Y, Liu M, Liu Y, Han M, Xiong X, et al. The modification role and tumor association with a methyltransferase: KMT2C. Frontiers in Immunology. 2024;15. 10.3389/fimmu.2024.1444923

41. Ni C, Wang X, Liu S, Zhang J, Luo Z, Xu B. KMT2C mutation as a predictor of immunotherapeutic efficacy in colorectal cancer. Scientific Reports. 2024;14:8284. 10.1038/s41598-024-57519-8

42. Nam C, Huang G, Zheng Y, Zhao H, Pan Y, Hu B, et al. The MLL3/GRHL2 complex regulates malignant transformation and anti-tumor immunity in squamous cancer. Journal of Experimental Medicine. 2025;222:e20240758. 10.1084/jem.20240758

43. Liu X, Xu Y, Wang Y, Jiang L, Yu C, Wang Y, et al. The pan-cancer mutational landscape of MLL3 and its impact on prognosis and immunochemotherapy. Cell Reports. 2025;44:116548. 10.1016/j.celrep.2025.116548

44. Liu C, Jin Y, Zhang H, Yan J, Guo Y, Bao X, et al. Effects of KMT2D mutation and its exon 39 mutation on the immune microenvironment and drug sensitivity in colorectal adenocarcinoma. Heliyon. 2023;9:e13629. 10.1016/j.heliyon.2023.e13629

45. Liu R, Niu Y, Liu C, Zhang X, Zhang J, Shi M, et al. Association of KMT2C/D loss-of-function variants with response to immune checkpoint blockades in colorectal cancer. Cancer Science. 2023;114:1229–39. 10.1111/cas.15716

46. Tauriello DVF, Palomo-Ponce S, Stork D, Berenguer-Llergo A, Badia-Ramentol J, Iglesias M, et al. TGFbeta drives immune evasion in genetically reconstituted colon cancer metastasis. Nature. 2018;554:538–43. 10.1038/nature25492

47. Tauriello DVF, Sancho E, Batlle E. Overcoming TGFβ-mediated immune evasion in cancer. Nat Rev Cancer. 2022;22:25–44. 10.1038/s41568-021-00413-6

